# Cas9-mediated Genome Editing Reveals a Significant Contribution of Calcium Signaling Pathways to Anhydrobiosis in Pv11

**DOI:** 10.1101/2020.10.15.340281

**Authors:** Yugo Miyata, Hiroto Fuse, Shoko Tokumoto, Yusuke Hiki, Ruslan Deviatiiarov, Yuki Yoshida, Takahiro G. Yamada, Richard Cornette, Oleg Gusev, Elena Shagimardanova, Akira Funahashi, Takahiro Kikawada

## Abstract

Pv11 is an insect cell line established from the midge *Polypedilum vanderplanki* that exhibits an extreme desiccation tolerance known as anhydrobiosis. Pv11 has also an anhydrobiotic ability which is induced by trehalose treatment. Here we report the successful construction of the genome editing system for Pv11 cells and its application for identifying the signaling pathways in the anhydrobiosis. Using the Cas9-mediated gene knock-in system, we established GCaMP3-stably expressing Pv11 cells to monitor intracellular Ca^2+^ mobilization. Intriguingly, trehalose treatment evoked a transient increase of cytosolic Ca^2+^ concentration, and further experiments indicated the contribution of the calmodulin – calcineurin – NFAT pathway to the tolerance for trehalose treatment as well as the desiccation tolerance, while the calmodulin – calmodulin Kinase – CREB pathway conferred only the desiccation tolerance on Pv11 cells. Thus, our results show the critical contribution of the trehalose–induced Ca^2+^ surge to the anhydrobiosis and the temporal different roles of each signaling pathway.

## Introduction

Anhydrobiosis is a striking ability of some organisms to survive under extreme desiccation. To date, among known anhydrobiotic animals, the sleeping chironomid *Polypedilum vanderplanki* is the only species from which culturable cell lines are established^1,2^. The cell line Pv11, derived from embryonic mass of the midge, recapitulate extreme desiccation tolerance of *P. vanderplanki* larvae^3^. Thus, Pv11 cells are a promising model for investigating the molecular mechanisms of the anhydrobiosis in *P. vanderplanki.* However, only a limited number of gene engineering tools were available in Pv11 cells^4,5^, and it has been a main experimental limitation in Pv11 cells.

Larvae of *P. vanderplanki* inhabit temporary rock pools in semi-arid regions of sub-Sahara in Africa^6^. During the drought season, the water in the rock pools dries up thoroughly and the larvae are also dehydrated. After sensing desiccation stimuli, the larvae start to synthesize excess of trehalose in the fat body tissues and deliver the trehalose to the whole body in order to protect biological molecules from desiccation damage by replacing intracellular water with a biological glass^6^. In addition to the physiological studies, genomics and metabolomics approaches have revealed the genes and metabolism related to the anhydrobiosis^6,7^.

To induce anhydrobiosis in Pv11 cells, high concentration trehalose pre-treatment is necessary^3^. During the pre-treatment, Pv11 cells uptake trehalose for the protection of biological molecules against forthcoming desiccation. In addition, trehalose treatment leads to changes in the gene expression profile^8,9^, suggesting that trehalose should be also a signal trigger for the gene expression, but exact mechanism of such interaction is unknown.

The CRISPR/Cas9 system is a versatile technology for genome editing due to its ease of use and accuracy and its conserved function in a wide range of species. Basically, the system is composed of two molecules, Cas9 nuclease and guide RNA (gRNA), and the complex makes a double strand break (DSB) at a target DNA site. When DNA fragments with the homology sequences around the DSB site exist, the exogenous DNA is efficiently inserted into the DSB site by intrinsic DNA repair mechanisms^10,11^. The knock-in technology has been used in various species including non-model organisms^12–15^. Therefore, the CRISPR/Cas9-based genome editing system for Pv11 cells should expand the experimental feasibility of exogenous gene expression and lead to revealing the detailed molecular mechanisms of anhydrobiosis in *P. vanderplanki*.

Genetically encoded fluorescent sensors are widely used to visualize molecular events and the mobilization of signaling molecules in living cells^16^. Generally, the fluorescent sensors are a fused protein that is composed of a sensing element and a reporting one, and the sensing element usually comprises an endogenous protein which binds to a specific molecule. The first developed class of fluorescent sensors is genetically encoded calcium indicators (GECIs) that can monitor intracellular Ca^2+^ levels^17^, and their sensitivity, selectivity and optical properties have been improved by many researchers^18^. GECIs have been used for investigating biological factors or molecules which increase intracellular Ca^2+^ concentrations and activate Ca^2+^ signaling pathways^19–23^, and using GECIs enables real-time measurement of Ca^2+^ signals in living cells. Thus, GECIs are useful to noninvasively examine if Ca^2+^ signals are related to a cellular function and when Ca^2+^ signals are evoked in living cells.

Ca^2+^ is an intracellular second messenger and the signaling pathways regulate large number of biological processes by regulating gene expression^24,25^. The key signaling pathways are calmodulin (CaM) – calcineurin (CaN) – nuclear factor of activated T-cells (NFAT) and CaM – calmodulin kinase (CaMK) – calcium/cAMP response element binding protein (CREB). CaM is a Ca^2+^ binding protein and transduces the change of intracellular Ca^2+^ concentration to molecular signals. CaN is a phosphatase activated by the Ca^2+^/CaM complexes and then dephosphorylates NFAT. Activated NFAT binds to the consensus sequence, TTTCCA, and regulates transcription of the downstream genes^26^. Ca^2+^/CaM complex-activated CaMK phosphorylates CREB, and it binds to the canonical CRE sequence, TGACGT, and induces the downstream gene expression. The signaling pathways are conserved in many cell types^24,27–29^ and across species^30–33^, and activated in response to several stresses, including oxidative stress^34,35^, heat shock^36^, osmotic stress^37^, and ER stress^38,39^. Therefore, we hypothesized that the Ca^2+^ signaling pathways can contribute to the induction of anhydrobiosis in *P. vanderplanki*.

Here we describe the construction and adaptation of the CRISPR/Cas9-based genome editing system for Pv11 cells and its application for identifying the signaling pathways required for the anhydrobiosis. Using the Cas9-mediated gene knock-in system, exogenous genes were inserted into the target site and constitutively expressed without affecting the anhydrobiosis. We applied the gene knock-in method to visualizing the mobilization of the second messenger, Ca^2+^, by knocking in the GCaMP3 gene that is one of GECIs. Amazingly, trehalose treatment evoked a transient increase of cytosolic Ca^2+^ concentration in Pv11 cells, and inhibition of Ca^2+^ signaling pathways during trehalose treatment decreased the survival rate after rehydration. The inhibitor experiment further indicated that the CaM – CaN – NFAT pathway conferred the tolerance for trehalose treatment on Pv11 cells as well as the desiccation tolerance, while the CaM - CaMK - CREB pathway conferred only the desiccation tolerance. These results show the critical contribution of the trehalose-induced Ca^2+^ surge to the anhydrobiosis and the distinct roles of signaling pathways.

## Results

### Utility of the CRISPR/Cas9 system in Pv11 cells

First, to examine if the CRISPR/Cas9 system works in Pv11 cells, Cas9-expression vector and synthetic gRNAs against AcGFP1 genes were transfected into Pv11-KH cells that stably express AcGFP1 (Fig. S1). After four weeks, the images of the cells were acquired (Fig. S1a), and flow cytometric analysis was carried out, showing that AcGFP1 fluorescence was disappeared in about 30% of the cells (Fig. S1b). The GFP-negative cells were sorted out, and the sequence of AcGFP1 gene on the genome was analyze. As shown in Figures S1c and d, insertion or deletion (indel) mutations were found both in gRNA#1- and #3-transfected cells (Figs. S1c and d). Together, these data showed that the CRISPR/Cas9 system was effectively workable to undergo targeted indel mutations in Pv11 cells.

### Construction of gRNA-expression vectors for Pv11 cells and the utility

To construct gRNA-expression vectors for Pv11 cells, the promoter region of the PvU6b gene was cloned (Supplementary Data 1). To overcome the experimental limitation that U6 promoters generally start the transcription with G or A^40–42^, DmtRNA sequence was added at the +1 position of the PvU6b promoter as described in the tRNA-flanked gRNA expression system^43–44^ (Supplementary Data 1). The gRNA- and Cas9-expression vectors were transfected into Pv11-KH cells, and after four weeks, the cell images and the flow cytometric data were acquired (Figs. S2a and b). As in the case of the synthetic gRNAs (Fig. S2), AcGFP1 fluorescence was disappeared in about 30% of the cells transfected both with Cas9- and gRNA-expression vectors (Fig. S2b). As shown in Figures S2c and d, indel mutations were found in the cells (Figs. S2c and d).

### CRISPR/Cas9-mediated targeted knock-in for exogenous gene expression

Targeted knock-in for exogenous gene expression is one of the most useful applications in the CRISPR/Cas9 system. To develop the knock-in technique in Pv11 cells, we tried the PITCh (Precise Integration of Target Chromosome) method^10^ and selected the *Pv.00443* gene as the insertion site because its expression is constitutively high under the regulation of a strong constitutive promoter, the 121 promoter, in Pv11 cells^5^. To exploit the endogenous constitutive expression system, we designed the gRNA targeting the 5′-flanking site of stop codon of *Pv.00443,* and then constructed the donor vector for the polycistronic expression of *Pv.00443,* AcGFP1 and ZeoR (Fig. 1a and Supplementary Data 2).

**Figure 1.**
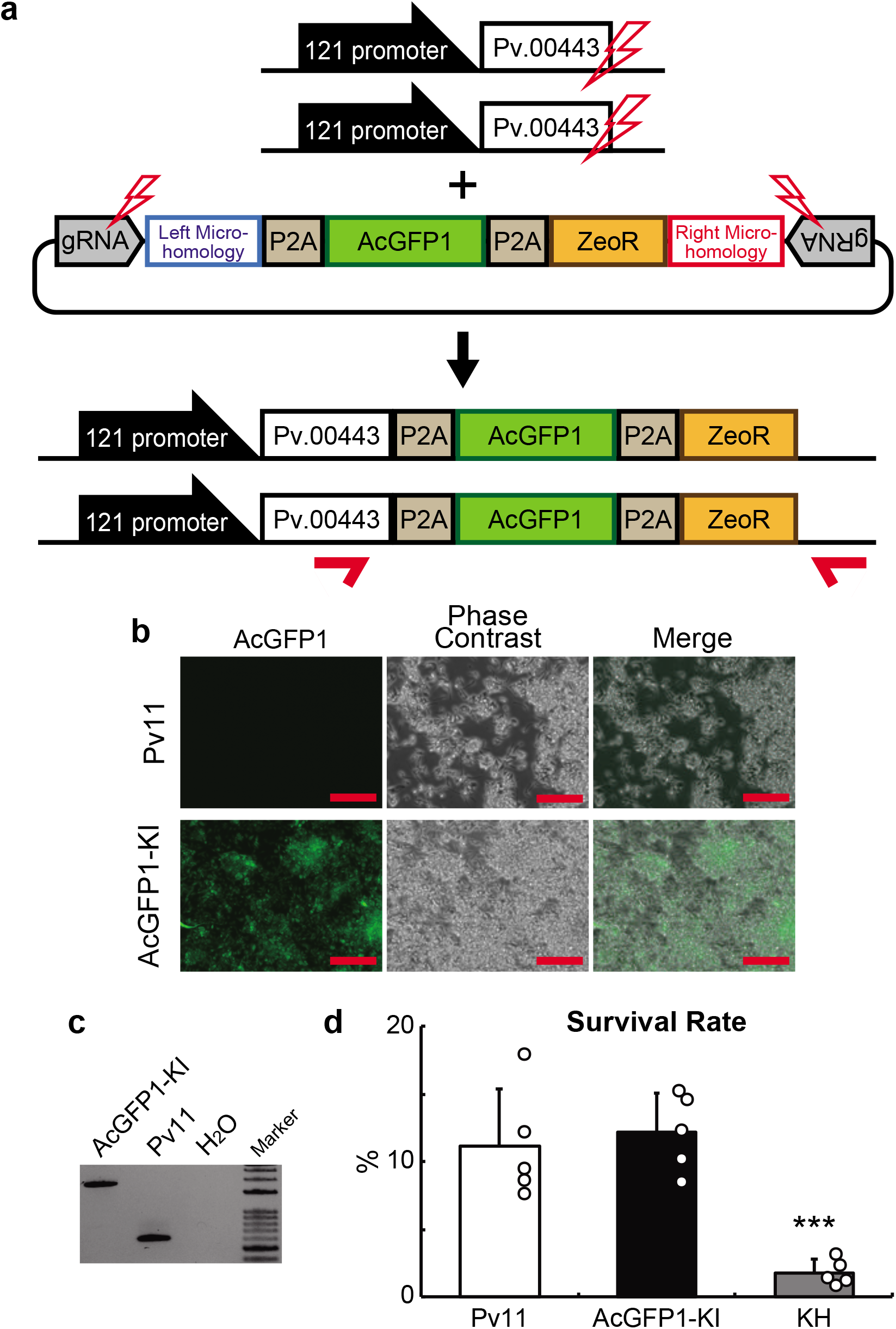
The AcGFP1 and zeocin resistance (ZeoR) genes were knocked in into 5′ flanking site of the stop codon of *Pv.00443* gene. **a)** The scheme of PITCh for AcGFP1- and ZeoR-knock-in is shown; the Cas9- and gRNA-expression vectors and the donor vector harboring AcGFP1 and ZeoR genes were transfected into Pv11 cells. Red arrows indicate the primers used in genomic PCR from the cells after selection by zeocin treatment and AcGFP1 fluorescence. **b)** The images of the cells were acquired by a microscope. **c)** Genomic PCR in the cells was carried out (the whole gel image is in Supplementary Fig. S3). **d)** The survival rate after desiccation-rehydration treatment was compared between two AcGFP1-stably expressing cells; targeted knock-in (AcGFP1-KI) and random integration-based (Pv11-KH) cells. An endogenous strong promoter, the 121 promoter, is located on the upstream region of the *Pv.00443* gene. Scale bars, 100 μm. The values are expressed as mean ± SD; n = 5 in each group. ***p = 0.0008 (vs. Pv11) and 0.0003 (vs. AcGFP1-KI).

The Cas9- and gRNA-expression vectors plus the donor vector were transfected into Pv11 cells, and the cells were treated with zeocin. After zeocin selection, single-cell sorting was performed by AcGFP1 fluorescence, and the sorted cell was proliferated with intact Pv11 cells as a feeder for two weeks. Zeocin treatment and sorting were carried out again to remove the feeder cells (Figs. 3b). The genome of the cells was extracted and subjected to genomic PCR. As shown in Figures 1c and S3, only a single 1840 bp-band was detected, which means biallelic integration with the donor construct (Figs. 1a, c and S3). Furthermore, the sequencing analysis of the band confirmed precise gene knock-in in Pv11 cells (Supplementary Data 3), and the established cell line was named AcGFP1-KI.

To assess an advantage of the targeted knock-in system over the random integrations, the survival rate after a desiccation-rehydration cycle was compared in two different AcGFP1-expressing cell lines, AcGFP1-KI and Pv11-KH; the latter was established by the random integrations of AcGFP1-expression units with the *PvGapdh* promoter^4^. As shown in Figure 1d, there was no significant difference in the survival rate between intact Pv11 and AcGFP1-KI cells, while the survival rate of Pv11-KH cells was significantly lower than the other two cell lines (p = 0.0008 for Pv11 and 0.0003 for AcGFP1-KI; Fig. 1d). This result clearly shows that the target knock-in into the 5′-flanking site of the stop codon of *Pv.00443* caused no deleterious effect on the anhydrobiotic ability of Pv11 cells.

To test if different donor DNAs can be inserted into the allele, two donor vectors were used (Fig. S4). The HaloTag and hygromycin resistance (HygR) genes were individually assigned on a donor vector and transfected with the Cas9- and gRNA-expression vectors (Fig. S4a). After hygromycin treatment, single-cell sorting was performed by staining with HaloTag fluorescent ligand, and the cell was proliferated with intact Pv11 cells as a feeder for two weeks. Hygromycin treatment and sorting were carried out again to remove feeder cells, and the cell images were acquired (Fig. S4b). The genome of the cells was extracted and subjected to genomic PCR. As shown in Figure S4c, 1139bp- and 1280-bp bands were detected, suggesting the genomic insertion of the HaloTag and HygR genes, respectively (Fig. S4c). Furthermore, the sequencing analysis of the PCR bands confirmed precise gene knock-in in Pv11 cells (Supplementary Data 4 and 5). Those results indicate the successful knock-in of two different donor DNAs into the same allele of homologous chromosomes and also the achievement of enough protein expression from a single copy of exogenous genes.

### Establishment of GCaMP3-KI cells

To apply the genome editing technique to reveal the signaling mechanisms in the anhydrobiosis desiccation tolerance, we tried to knock-in the GCaMP3 gene, which is one of GECIs used in a wide range of cell types^20^, including insect cells^45,46^. As shown in Figure 2a, the Cas9- and gRNA-expression vectors plus two different donor vectors harboring GCaMP3 and ZeoR were transfected into Pv11 cells (Fig. 2a). To select the cells expressing both genes, zeocin treatment and subsequent single cell sorting were performed (Fig. S5). The clonal cell line exhibited strong GCaMP3 fluorescence after treated with the calcium ionophore, ionomycin, compared to DMSO (Fig. 2b). The genome of the cells was extracted, and subsequent genomic PCR showed 2035bp- and 1057bp-bands, suggesting precise GCaMP3- and ZeoR-knock-in, respectively (Figs. 2c and S6). Sequencing analysis of the PCR bands confirmed precise gene knock-in in Pv11 cells (Supplementary Data 6 and 7). Furthermore, the desiccation tolerant ability of the KI cells was the same with Pv11 cells (Fig. 2d). We named the stably GCaMP3-expressing cells GCaMP3-KI.

**Figure 2.**
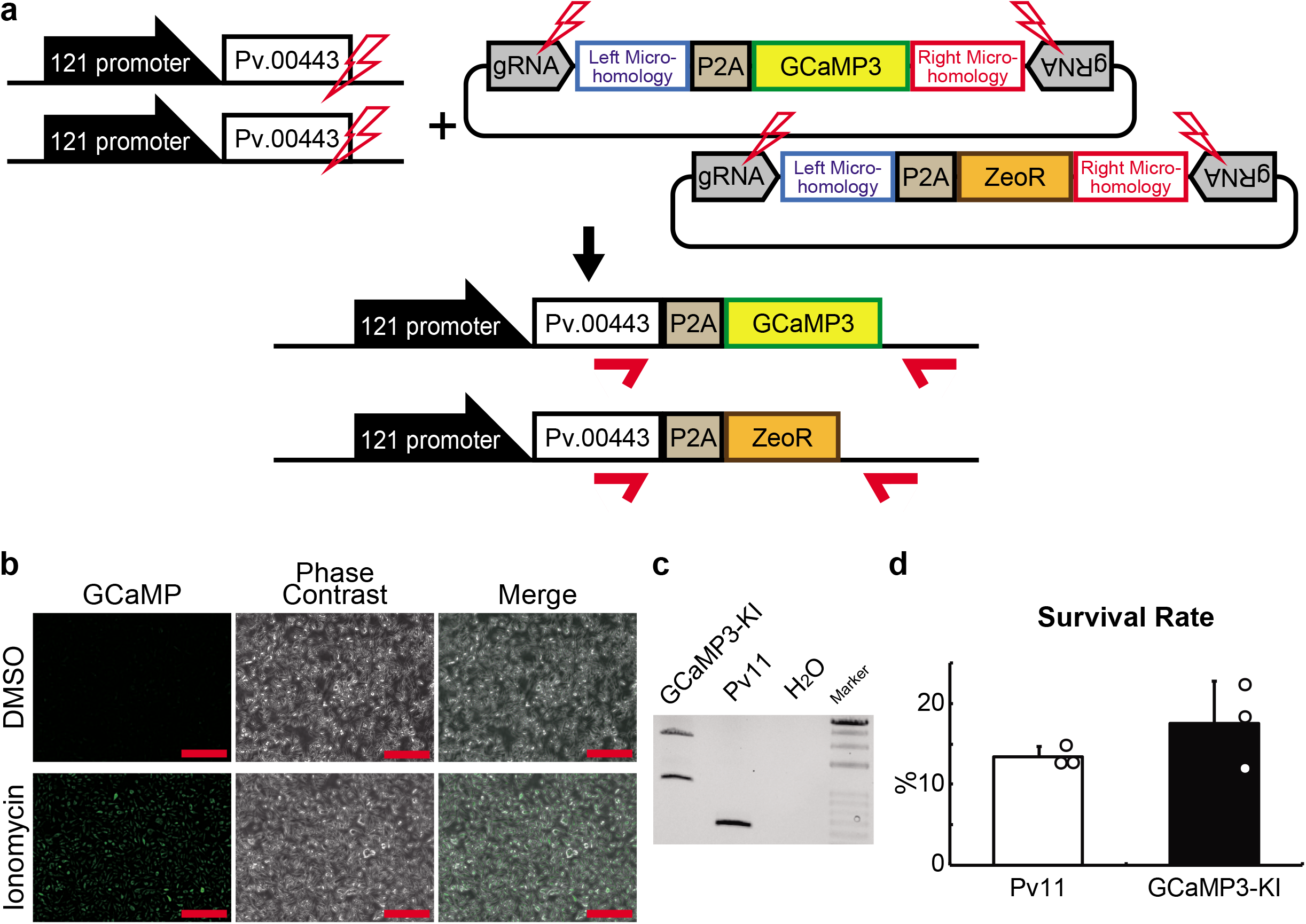
The establishment of a stably GCaMP3-expressing cell line. **a)** The scheme of PITCh for GCaMP3- and ZeoR-knock-in is shown; the Cas9- and gRNA-expression vectors and two donor vectors harboring GCaMP3 or ZeoR gene were transfected into Pv11 cells. Red arrows indicate the primers used in genomic PCR from the cells after selection by zeocin treatment and GCaMP3 fluorescence. **b)** The images of the cells treated with DMSO or Ionomycin were acquired by a microscope. **c)** Genomic PCR in the cells was carried out (the whole gel image is in Supplementary Fig. S6) **d)** The survival rate after desiccation-rehydration treatment was compared between Pv11 and GCaMP3-KI cells. Scale bars, 100 μm. The values are expressed as mean ± SD; n = 3 in each group.

**Figure 3.**
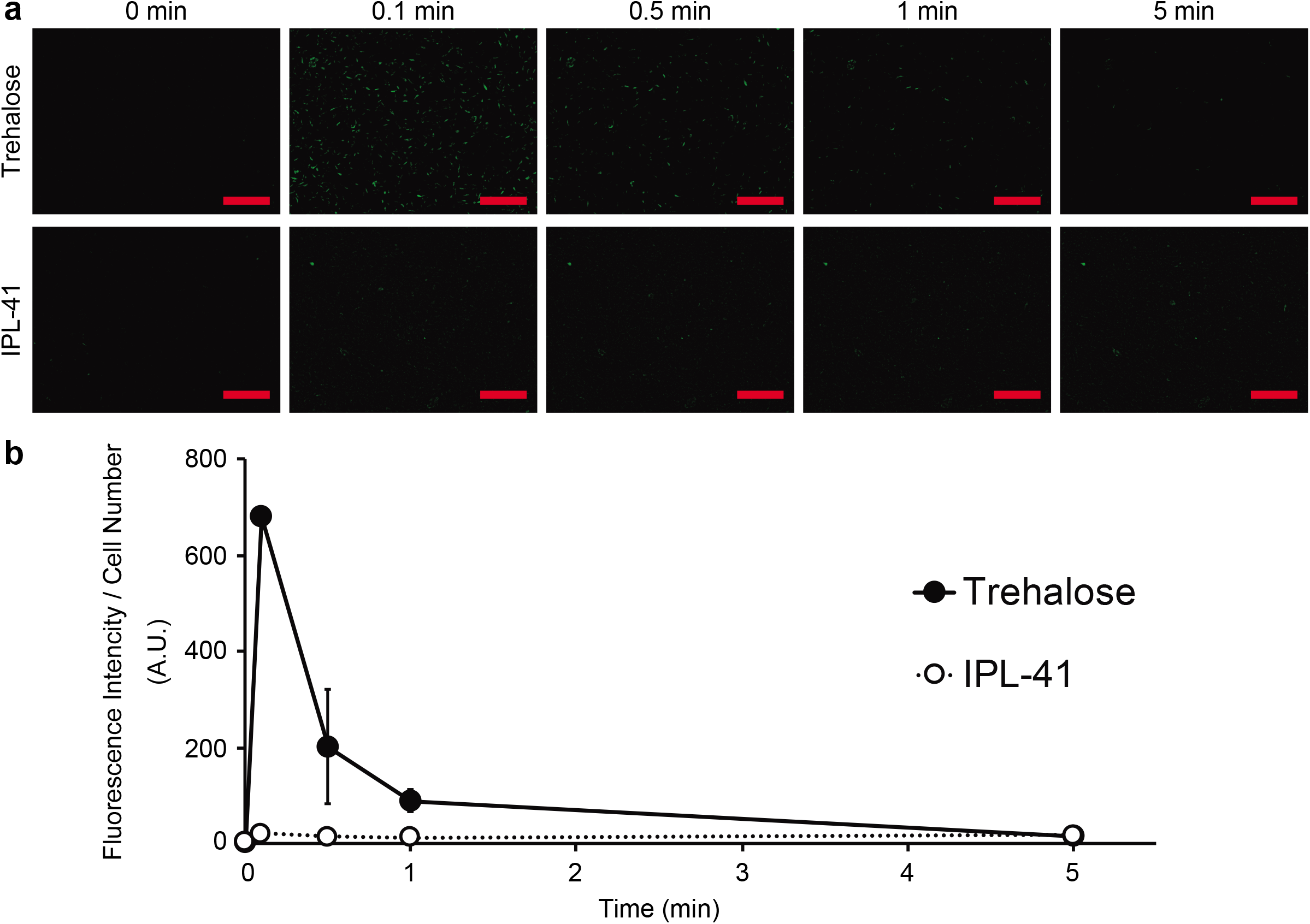
Intracellular Ca^2+^ mobilization of GCaMP3-KI cells during trehalose treatment. **a)** Trehalose treatment induced a transient increase of GCaMP3-fluorescence in the cells. **b)** Quantitative data were shown. Scale bars, 100 μm. The values are expressed as mean ± SD; n = 3 in each group.

### Intracellular Ca^2+^ levels during trehalose treatment

We examined whether the intracellular Ca^2+^ concentration changed in Pv11 cells during trehalose treatment as the expression pattern of the desiccation-related genes was altered^8^. GCaMP3-KI cells were treated with trehalose, and the fluorescence was monitored. As shown in Figures 3a, S7 and S8, trehalose treatment induced a transient increase of GCaMP3- fluorescence in GCaMP3-KI cells (Figs. 3a, S7 and S8), while ionomycin treatment stably increased the GCaMP3 fluorescence (Fig. S9). Quantitative analysis of GCaMP3 fluorescence showed 236-fold increase at 0.1min after trehalose treatment, and the fluorescent intensity returned to the basal level within five minutes (Figs. 3b and S10).

### Contribution of calcium signaling pathways to the desiccation tolerance

To examine the contribution of the cytosolic calcium surge on the desiccation tolerance, Pv11 cells were treated with inhibitors against calcium signaling pathways during desiccation-rehydration processes (Fig. S11a). At first, we examined the inhibitor concentrations at which no deleterious effect on the survival rate in normal culture media, IPL-41 (Figs. S11b, c, and 4a-g, and Supplementary Table 1). At the same concentrations, the CaM inhibitors (W7 and Calmidazolium), the CaN inhibitors (FK-506 and Cyclosporin A) and the NFAT inhibitor (INCA-6)^47^ decreased the survival rate after 48h-incubation of trehalose (Figs. 4a-c and f), while the CaMK inhibitors (KN-93 and STO-609) and the CREB inhibitor (666-15)^48^ did not have such effect (Figs. 4d, e and g). All the inhibitors decreased the survival rate after rehydration (Figs. 4a-g). These results demonstrate the following two things; i) the activation of the calcium signaling pathways should be necessary for desiccation tolerance of Pv11 cells, ii) the CaM - CaN - NFAT pathway confers the tolerance for trehalose treatment on Pv11 cells as well as the desiccation tolerance.

**Figure 4.**
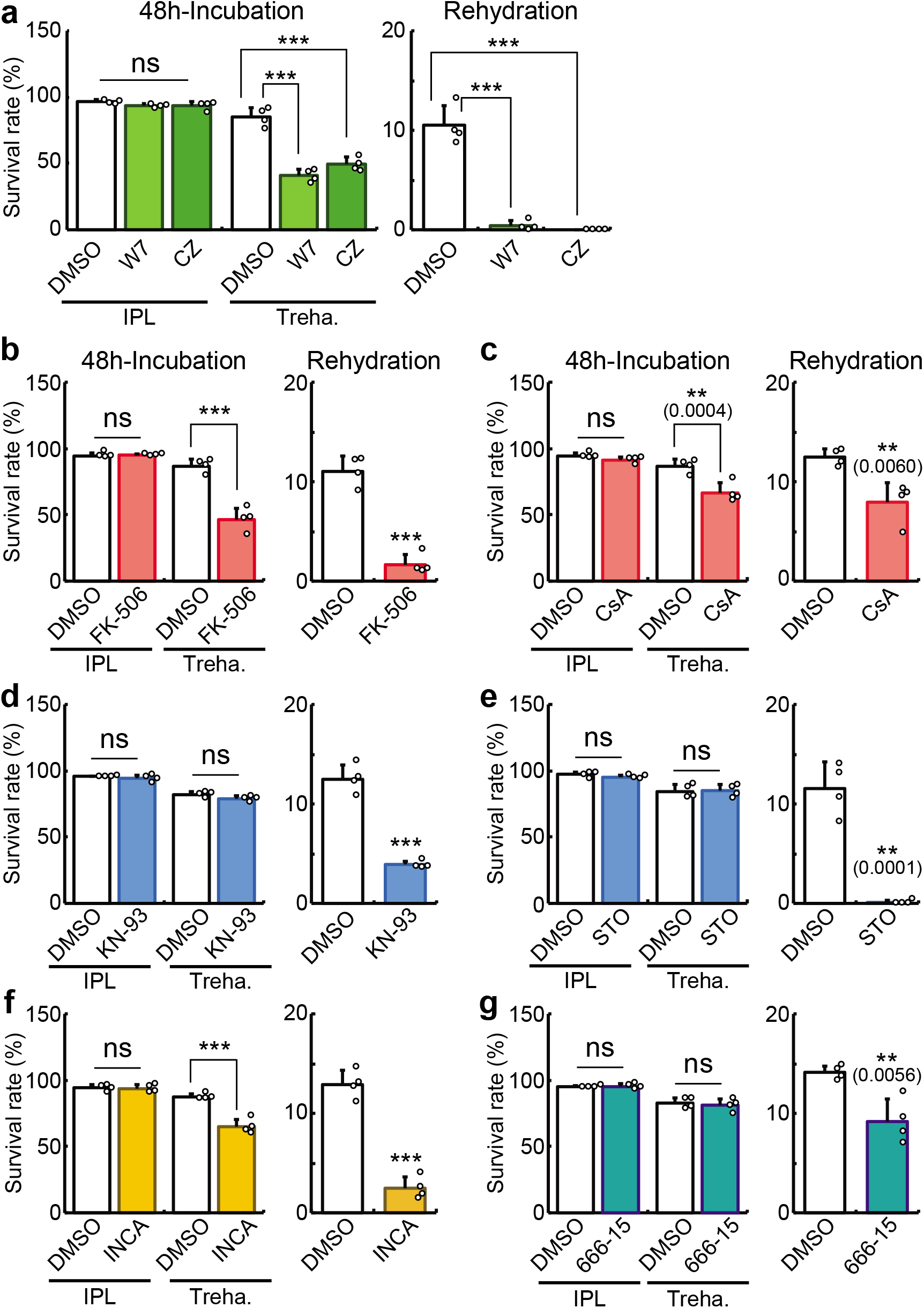
Contribution of the calcium signaling pathways to the desiccation tolerance **a-g)** CaM inhibitors (W7 and Calmidazolium; a), CaN inhibitors (FK-506 and Cyclosporine A; b and c, respectively), CaMK inhibitors (KN-93 and STO-609; d and e, respectively), an NFAT inhibitor (INCA-6; f) and a CREB inhibitor (666-15; g) were treated in Pv11 cells and the survival rates were analyzed during IPL-41/trehalose incubation or after desiccation-rehydration treatment. The values are expressed as mean ± SD; n = 4 in each group. Each inhibitor was treated at 100μM of W7, 7.5μM of Calmidazolium, 40μM of FK-506, 160nM of Cyclosporin A, 30μM of KN-93, 20μM of STO-609, 30μM of INCA-6, and 20μM of 666-15. ***p < 0.0001, **p < 0.01 (actual p-values are shown within parentheses).

## Discussion

The major achievements of the current study are: i) successful construction of genome editing tools for Pv11 cells; and ii) the application of the technique for revealing the molecular signalings responsible for anhydrobiosis in Pv11 cells. The Cas9-mediated genome editing technique has been improved and now is a basic tool for current molecular biology even in non-model species^49-52^. Here we used the CRIS-PITCh method for inserting genes of interests (Figs. 1, 2 and S4), and one of the KI cell lines, GCaMP3-KI, uncovered a novel function of trehalose in evoking an intracellular calcium surge and the contribution of the calcium signaling pathways to the anhydrobiosis. These results suggested a critical role of trehalose as a signal inducer.

The CRISPR/Cas9 system was functionally available in Pv11 cells with synthetic gRNAs or the gRNA expression vectors (Figs. S1 and S2). Furthermore, targeted knock-in was achieved using the PITCh method (Figs. 1, 2, and S4), and it enabled the establishment of stably expressing cell lines without losing the anhydrobiotic ability (Figs. 1d, 2d, and S4d). In our forthcoming study, we plan to disrupt an endogenous gene and are now constructing the donor vectors for creating knock-in/knock-out alleles^10^. Thus, the current success of targeted knock-in in Pv11 cells leads to endogenous gene knock-out and thereby to identify the genes essential for the anhydrobiosis.

To improve our understanding of the molecular mechanisms underlying the anhydrobiosis in Pv11 cells, we have developed several gene manipulation methods, especially in exogenous gene expression using plasmid vectors^4,5,53^. However, the transfection method for Pv11 cells is limited to electroporation^4^, and the transfection damage decrease desiccation tolerance of Pv11 in transient expression experiments, likely due to temporary damages by the electroporation^54,55^ (Fig. S12). To avoid the temporary damages, we generated stably AcGFP1-expressing Pv11 cells (Pv11-KH cells) through random integration of an expression vector, but the cells showed lower desiccation tolerance than intact Pv11 cells^4^, maybe due to the unintended disruption of desiccation-related genes by genomic insertion of the exogenous DNA fragments^56,57^. Thus, to express exogenous genes with no deleterious effects on the desiccation tolerance, a targeted insertion system is necessary in Pv11 cells.

GCaMP3-KI cells showed strong fluorescence immediately after trehalose treatment (Fig. 3), suggesting that Pv11 cells sense a specific factor or factors from trehalose. What does transduce extracellular trehalose into the calcium signal? We hypothesize that osmotic sensors and mechanosensors, such as OSCA, Piezo and TRP cation channels^58–60^, can be responsible for the Ca^2+^ mobilization. Gating the channels allow mono- or di-valent cations such as Na+ or Ca^2+^to enter inside the cells, potentially activating calcium signaling pathways. To examine the contribution to the calcium mobilization, we are currently constructing the knock-out cell lines. Another candidate is a gustatory receptor that senses trehalose, like Gr5a^61,62^. In Gr5a-expressing *Drosophila* S2 cells, trehalose induced an intracellular calcium surge, suggesting that the receptor functioned in multiple cellular types. Furthermore, recent researches showed the possibility that several gustatory receptors expressed in non-gustatory receptor-expressing neurons have different roles of canonical gustatory receptors apart from chemosensation^63,64^. To examine the possibility, *in silico* screening and experimental confirmation may be needed^65,66^. We believe that these approaches will clarify the molecular mechanism of the trehalose sensing and the signaling.

Our result clearly showed the critical role of calcium signaling pathways in anhydrobiosis in Pv11 cells (Fig. 4). The potential contribution of calcium signalings to anhydrobiosis was also suggested in tardigrades and nematodes^67,68^. In the anhydrobiotic tardigrade, *Hypsibius dujardini,* a calmodulin inhibitor (J-8) and a calcium release modulator (2-APB) reduced the survival rate after rehydration, but the inhibitors against CaN and CaMK had no effect on the anhydrobiotic ability. Given that calmodulin regulates the gating of calcium ion channel in endoplasmic reticulum and mitochondrion as well as the signaling pathways^69,70^, the J-8 and 2-APB treatments may disrupt calcium storage in the organelles rather than calcium signalings, leading to malfunction of those intracellular organelles^71,72^. RNAi-based screening using the nematode *Panagrolaimus superbus,* showed that knockdown of “CAMK/CAMKL/MELK protein kinase” decreased the survival rate after rehydration^68^, whereas the contribution of CaN - NFAT pathway or CREB on the anhydrobiotic ability was not assessed. Our detailed experiments show that the CaM - CaN - NFAT pathway activation confers the tolerance to both trehalose incubation and desiccation on Pv11 cells, while the CaM - CaMK - CREB pathway activation gives only desiccation tolerance (Figs. 4 and S11). Therefore, the differential functions of the individual calcium pathway in anhydrobiosis are shown for the first time in our study.

In the current study, we succeeded in detecting Ca^2+^ mobilization using GCaMP3-KI cells. There are a lot of second messengers reported, and the knock-in technique can be applied to them since a wide variety of genetically encoded fluorescent sensors have been developed^16^. For example, cyclic AMP, inositol 1,4,5-trisphosphate, diacylglycerol and reactive oxygen species^73,74^ can be detected by their specific fluorescent sensors. In addition, the active status of proteases, kinases, and GTPases^75,76^ can be visualized. Thus, real-time monitoring of these biomolecules should uncover the anhydrobiosis-specific signaling profile. Another advantage of genetically encoded fluorescence sensors is controllability of their subcellular localization^75,77^. Given that differential subcellular localization of channels and enzymes contributes to their specific functions^78^–^84^, analyzing the active state of signal pathways with subcellular resolution should be helpful for comprehensive understanding of the anhydrobiosis-related signaling profile.

In conclusion, genome editing with the CRISPR/Cas9 system was successfully developed in Pv11 cells, and it revealed the critical contribution of the calcium signaling pathways to the anhydrobiosis. The technique will allow further advanced molecular engineering, for example, gene knockout, and enable us to characterize the mechanisms underlying anhydrobiosis in Pv11 cells.

## Methods

### Cell culture

Pv11 and Pv11-KH cells were cultured in IPL-41 medium (Thermo Fisher Scientific) supplemented with 2.6 g/L tryptose phosphate broth (Becton, Dickinson and Company, Franklin Lakes, NJ, USA), 10% (v/v) fetal bovine serum, and 0.05% (v/v) of an antibiotic and antimycotic mixture (penicillin, amphotericin B, and streptomycin; Sigma Aldrich, St. Louis, MO, USA)^5^, designated hereafter as complete IPL-41 medium.

### Expression vectors

The SpCas9-expressing vector was constructed by replacing the AcGFP1 region of pPv121-AcGFP1, using a HiFi Assembly kit (New England BioLabs, Ipswich, MA)^53^. The SpCas9 gene was cloned from pBS-Hsp70-Cas9, which was a gift from Melissa Harrison & Kate O’Connor-Giles & Jill Wildonger (Addgene plasmid # 46294; http://n2t.net/addgene:46294; RRID: Addgene_46294).

The PvU6b promoter was cloned after reference to the gene model in MidgeBase (http://bertone.nises-f.affrc.go.jp/midgebase/). The pPvU6b plasmid was constructed by replacing DmU6 promoter of pU6-BbsI-chiRNA with the PvU6b promoter and inserting DmtRNA sequence described in the previous articles^43,44^. pU6-BbsI-chiRNA was a gift from Melissa Harrison & Kate O’Connor-Giles & Jill Wildonger (Addgene plasmid # 45946)^85^, and we name the vector pPvU6b-DmtRNA-BbsI (the complete sequence is shown in Supplementary Data 1). To express the gRNA targeting the GFP gene or the 5′-flanking site of the stop codon of *Pv.00443* (Figs. S2 and 1), pPvU6b-DmtRNA-Pv.00443#1 was constructed as below; i) pPvU6b-DmtRNA-BbsI was restricted with BbsI, ii) the pair of two oligo DNAs were annealed (Supplementary Table 2), and iii) the annealed DNA was ligated into the cut vector (the complete sequences of the gRNA-expression vectors are shown in Supplementary Data 8, 9 and 10).

### Donor vectors

The donor vector having the AcGFP1-P2A-ZeoR sequence (Fig. 1) was constructed using PCR, a HiFi Assembly kit (New England BioLabs) and a TOPO cloning kit (Thermo Fisher Scientific, Waltham, MA, USA). Briefly, at first, the polycistronic sequence of AcGFP1-P2A-ZeoR was constructed on the pPv121 vector, using a HiFi Assembly kit (New England BioLabs). Then, the vector was used as a PCR template to add another P2A sequence to the 5′ end of AcGFP1, and the PCR product was inserted into pCR-Blunt II-TOPO vector (Thermo Fisher Scientific). Furthermore, the vector was used as a PCR template to add gRNA-target and microhomology sequences, and the PCR product was also cloned into pCR-Blunt II-TOPO vector (the complete sequence is shown in Supplementary Data 2). To construct the donor vector for inserting the HaloTag gene (Fig. S4), pCR4 Blunt-TOPO vector was used for cloning. To construct the donor vector for inserting the other genes, we created the basic vector, pCR4-Pv.00443#1μH-P2A-BbsI, which has the sequences of the gRNA-target, microhomology, P2A, and BbsI; at first, synthetic gene with the sequences was acquired for the PCR template (eurofins Genomics, Tokyo, Japan), and the PCR product was cloned into pCR4 Blunt-TOPO vector (the complete sequence is shown in Supplementary Data 11). To construct the donor vectors for inserting BlaR, ZeoR, and GCaMP3 genes (Figs. S4 and 2), the basic vector was restricted by BbsI (New England BioLabs) and HiFi Assembly was performed. The HaloTag, BlaR and GCaMP3 genes were cloned from pFN19K HaloTag T7 SP6 (Promega, Fitchburg, WI, USA), pYES6/CT (Thermo Fisher Scientific) and GCaMP (Plasmid #22692, Addgene) (doi.org/10.1038/nmeth.1398) (the complete sequence is shown in Supplementary Data 12, 13, 14 and 15).

### Transfection, drug selection and cell sorting

The cells used in each experiment were seeded at a density of 3 × 10^5^ cells per mL into a 25 cm^2^ cell culture flask and grown at 25°C for 4-6 days before transfection. Transfection was carried out using a NEPA21 Super Electroporator (Nepa Gene, Chiba, Japan) as described previously^4^. The 5 μg of gRNA- and SpCas9-expression vectors plus 0.03-0.1 pmol of donor vectors were transfected into the cells. Five days after transfection, the cells were treated with 400 μg/mL zeocin or 200 μg/mL blasticidin at a density of 1 × 10^5^ cells per mL. After the one-week drug selection, the medium was changed to normal IPL-41 medium, then the cells were cultured for two weeks. To establish the clonal cell lines, single cell sorting was performed by a MoFlo Astrios cell-sorter equipped with 355-, 488- and 640-nm lasers. One thousand intact Pv11 cells were seeded as a feeder on a well of a 96-well plate prior to the sorting. HaloTag-positive cells were labeled with Janelia Fluor 646 HaloTag fluorescent Ligand (Promega). GCaMP3-KI cells exhibited strong fluorescence when subjected to a flow cytometer maybe due to shear or/and mechanical stress (Fig. S5). The cells were stained with DAPI (Dojindo, Kumamoto, Japan), and the fluorescence of DAPI, GCaMP3 and Janelia Fluor 646 HaloTag fluorescent Ligand were excited with 355-nm, 488-nm and 640-nm lasers, respectively. After single cell sorting, the cells were cultured for 2 weeks, and zeocin or blasticidin was treated to eliminate feeder cells. If the drug selection was not enough, bulk sorting was performed.

### Genomic PCR and sequencing analysis

To confirm the precise gene knock-in, genomic PCR and sequencing analysis were performed (Figs. 1, S4 and 2). For genomic PCR, the genome of Pv11 cells and the clonal cell lines were extracted with a NucleoSpin Tissue kit (Takara Bio, Shiga, Japan) and subjected to PCR using the following primer set; 5′-GCCAAAGCGAGCCAATTCAA-3′ and 5′- GGGTGTTATTGCTACTTTAATGCGT-3′. The PCR images were acquired by a ChemiDoc Touch imaging system (Bio-Rad, Hercules, CA, USA). After gel purification of the PCR products, sub-cloning was carried out using a TOPO cloning kit (Thermo Fisher Scientific), and the plasmids were subjected to sequencing analysis.

### Desiccation and rehydration

Pv11 cells were subjected to desiccation-rehydration (Figs. 1d, S4d, 2d and 4) as described previously (Watanabe et al. 2016). Briefly, Cells were incubated in a preconditioning medium (600 mM trehalose containing 10% (v/v) complete IPL-41 medium) for 48 h at 25 ° C, and then suspended in 400 μL of the preconditioning medium. Forty microliter aliquots of the cell suspension were droped into 35-mm petri dishes, and the dishes were desiccated and maintained at < 10% relative humidity and 25 ° C for more than 10 days. An hour after rehydration by complete IPL-41 medium, cells were stained with Propidium iodide (PI; Dojindo) and Hoechst 33342 (Dojindo), and then the images were acquired by a BZ-X700 microscope (Keyence, Osaka, Japan). The survival rate was calculated by the ratio of the number of the live cells (Hoechst positive and PI negative) to that of total cells (Hoechst-positive).

### GCaMP3 fluorescence quantification

The images for GCaMP3 fluorescence quantification were acquired as follows; i) 50μl of complete IPL-41 media with 10μg/ml of Hoechst 33342 (Dojindo) was put on a well of 48-well plate, ii) 50μl of cell suspension was added on the well, and the cells were allowed to attach the culture surface (usually for 7 to 10 minutes), iii) a “0 min” image was acquired, iv) 900μl of 600mM trehalose, complete IPL-41 or complete PL-41 with ionomycin was added into the well, and time course images were acquired. The brightness of GCaMP fluorescence was quantified using Hybrid Cell Count BZ-H2C software (Keyence). The cell numbers were also counted from Hoechst 33342 fluorescence using the same software.

### Ionomycin or inhibitor treatment

GCaMP3-KI cells were treated with ionomycin at 10μM (Figs. 2b, S9 and S10, and Supplementary Table 1). To inhibit the calcium signalings by trehalose treatment, Pv11 cells were pretreated with the inhibitors for one hour, and then treated with trehalose or complete IPL-41 plus the inhibitors (Figs. 4 and S11b, and Supplementary Table 1).

### Statistics and reproducibility

All data were expressed as mean ± SD. Differences between two groups were examined for statistical significance by the Student t-test in Figs. 2d and 4. Differences among more than three groups were examined by ANOVA followed by a Tukey test (Figs. 1d and 4). A P-value < 0.05 denoted a statistically significant difference. GraphPad Prism 8 software (GraphPad, San Diego, CA) was used for the statistical analyses. Experiments were independently repeated at least three times in general. The raw data underlying plots in the figures are available in Supplementary Data 16.

## Supporting information

Description of Supplementary Data

## Data availability

The datasets shown in the current study are available from the corresponding author on reasonable request. The source data of Figures 1d, 2d, 3b and 4a-g, and Supplementary Figures S1b, S2b, S4d, S5b, S5c, S10, S11c, and S12 are provided in Supplementary Data 16.

## Acknowledgements

We would like to express our deep gratitude to Tomoe Shiratori (NARO) for their technical support.

This work was supported by Grants-in-Aid for Scientific Research (KAKENHI) Grants (numbers JP25128714& JP17H01511 to T.K., JP18K14472 to Y.M., JP19J12030 to S.T., JP18J21155 to Y.Y., JP16K07308 to R.C. and JP18H02217 to O.G.), and T.K. and R.C. were also funded by the Agriculture, Forestry and Fisheries Research Council of the Ministry of Agriculture, Forestry and Fisheries (www.affrc.maff.go.jp) grant “Pilot program of international collaborative research (Collaborative research based on a joint call with Russia)” under “Commissioned projects for promotion of strategic international collaborative research” (JPJ008837). E.S. and R.D. were supported by Russian Science Foundation grant 20-44-07002 for bioinformatics part of the study.

## Author contributions

Y.M. performed the experiments, analyzed the data, contributed to discussion, and wrote the manuscript. H.F. and S.T. performed the experiments, analyzed the data and contributed to discussion.

Y.H., Y.Y. and T. Y. performed the experiments, analyzed the data, contributed to discussion and wrote the manuscript.

R. D. analyzed the data and contributed to discussion.

R.C., A.F., O.G., and E.S. contributed to discussions and participated in editing the manuscript. T.K. designed the project, wrote the manuscript, and contributed to discussions.

