## Supplementary material for "Cas9-mediated Genome Editing Reveals a Significant Contribution of Calcium Signaling Pathways to Anhydrobiosis in Pv11": Supplemenatry Figure

Fig. S1

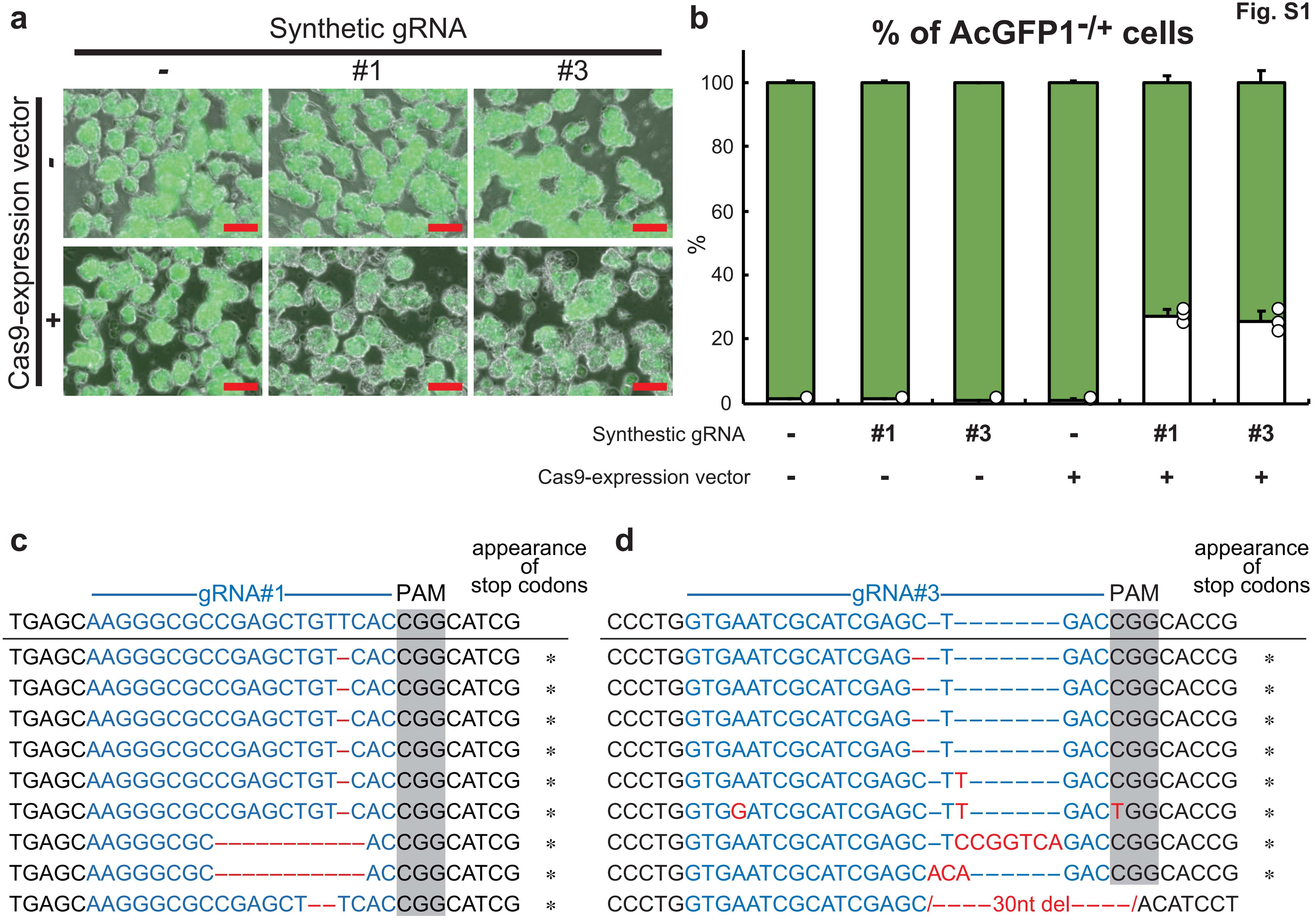

a
Synthetic gRNA
-
#1
#3
Cas9-expression vector
-
+
b
% of AcGFP1-/+ cells
100
80
60
40
20
0
%
Synthetic gRNA - #1 #3 - #1 #3
Cas9-expression vector - - - + + +
c
gRNA#1
PAM
appearance of stop codons
TGAGCAAGGGCGCCGAGCTGTTCACCGGCATCG
TGAGCAAGGGCGCCGAGCTGT-CACCGGCATCG *
TGAGCAAGGGCGCCGAGCTGT-CACCGGCATCG *
TGAGCAAGGGCGCCGAGCTGT-CACCGGCATCG *
TGAGCAAGGGCGCCGAGCTGT-CACCGGCATCG *
TGAGCAAGGGCGCCGAGCTGT-CACCGGCATCG *
TGAGCAAGGGCGCCGAGCTGT-CACCGGCATCG *
TGAGCAAGGGCGC----------ACCGGCATCG *
TGAGCAAGGGCGC----------ACCGGCATCG *
TGAGCAAGGGCGCCGAGCT--TCACCGGCATCG *
d
gRNA#3
PAM
appearance of stop codons
CCCTGGTGAATCGCATCGAGC-T-------GACCGGCACCG
CCCTGGTGAATCGCATCGAG--T-------GACCGGCACCG *
CCCTGGTGAATCGCATCGAG--T-------GACCGGCACCG *
CCCTGGTGAATCGCATCGAG--T-------GACCGGCACCG *
CCCTGGTGAATCGCATCGAG--T-------GACCGGCACCG *
CCCTGGTGAATCGCATCGAGC-TT------GACCGGCACCG *
CCCTGGTGGATCGCATCGAGC-TT------GACTGGCACCG *
CCCTGGTGAATCGCATCGAGC-TCCGGTCAGACCGGCACCG *
CCCTGGTGAATCGCATCGAGCACA------GACCGGCACCG *
CCCTGGTGAATCGCATCGAGC/----30nt del----/ACATCCT

Supplementary Figure S1

The CRISPR/Cas9 system using synthetic gRNAs in Pv11 cells.

**a)** The Cas9-expression vector and the synthetic gRNAs targeting the AcGFP1 gene were transfected into Pv11-KH cells, and the cells were grown for more than four weeks. **b)** The proportions of AcGFP1-negative/positive cells were analyzed by a flow cytometer. **c, d)** The genomic sequences were analyzed in the AcGFP1-negative cells which were transfected with synthetic gRNA #1 (c) or #3 (d). Scale bars, 100 μm. The values are expressed as mean ± SD; n = 3 in each group. Red letters and dashes indicate insertion and deletion mutations, respectively.

Fig. S2

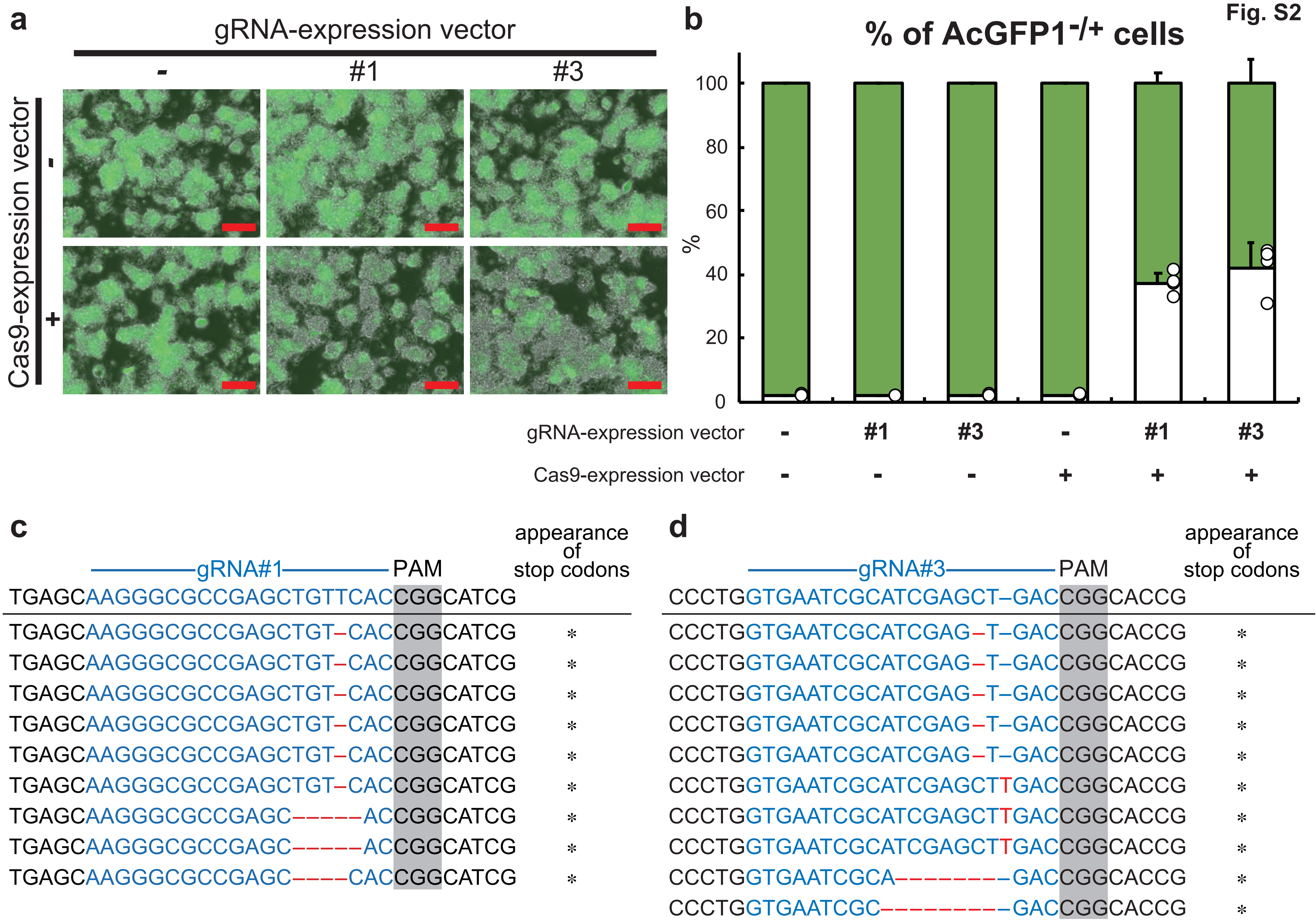
a
gRNA-expression vector
-
#1
#3
Cas9-expression vector
-
+
b
% of AcGFP1-/+ cells
100
80
60
40
20
0
%
gRNA-expression vector - #1 #3 - #1 #3
Cas9-expression vector - - - + + +
c
gRNA#1
PAM
appearance of stop codons
TGAGCAAGGGCGCCGAGCTGTTCACCGGCATCG
TGAGCAAGGGCGCCGAGCTGT–CACCGGCATCG *
TGAGCAAGGGCGCCGAGCTGT–CACCGGCATCG *
TGAGCAAGGGCGCCGAGCTGT–CACCGGCATCG *
TGAGCAAGGGCGCCGAGCTGT–CACCGGCATCG *
TGAGCAAGGGCGCCGAGCTGT–CACCGGCATCG *
TGAGCAAGGGCGCCGAGCTGT–CACCGGCATCG *
TGAGCAAGGGCGCCGAGC–––––ACCGGCATCG *
TGAGCAAGGGCGCCGAGC–––––ACCGGCATCG *
TGAGCAAGGGCGCCGAGC––––CACCGGCATCG *
d
gRNA#3
PAM
appearance of stop codons
CCCTGGTGAATCGCATCGAGCT–GACCGGCACCG
CCCTGGTGAATCGCATCGAG–T–GACCGGCACCG *
CCCTGGTGAATCGCATCGAG–T–GACCGGCACCG *
CCCTGGTGAATCGCATCGAG–T–GACCGGCACCG *
CCCTGGTGAATCGCATCGAG–T–GACCGGCACCG *
CCCTGGTGAATCGCATCGAG–T–GACCGGCACCG *
CCCTGGTGAATCGCATCGAGCTTGACCGGCACCG *
CCCTGGTGAATCGCATCGAGCTTGACCGGCACCG *
CCCTGGTGAATCGCATCGAGCTTGACCGGCACCG *
CCCTGGTGAATCGCA––––––––GACCGGCACCG *
CCCTGGTGAATCGC–––––––––GACCGGCACCG *

Supplementary Figure S2

The CRISPR/Cas9 system using gRNA-expression vectors in Pv11 cells.

**a)** The Cas9- and gRNA-expression vectors were transfected into Pv11-KH cells, and the cells were grown for more than four weeks. **b)** The proportions of AcGFP1-negative/positive cells were analyzed by a flow cytometer. **c, d)** The genomic sequences were analyzed in AcGFP1-negative cells which were transfected with #1- (c) or #3- (d) gRNA expression vectors. Scale bars, 100 μm. The values are expressed as mean ± SD; n = 4 in each group. Red letters and dashes indicate insertion and deletion mutations, respectively.

Fig. S3

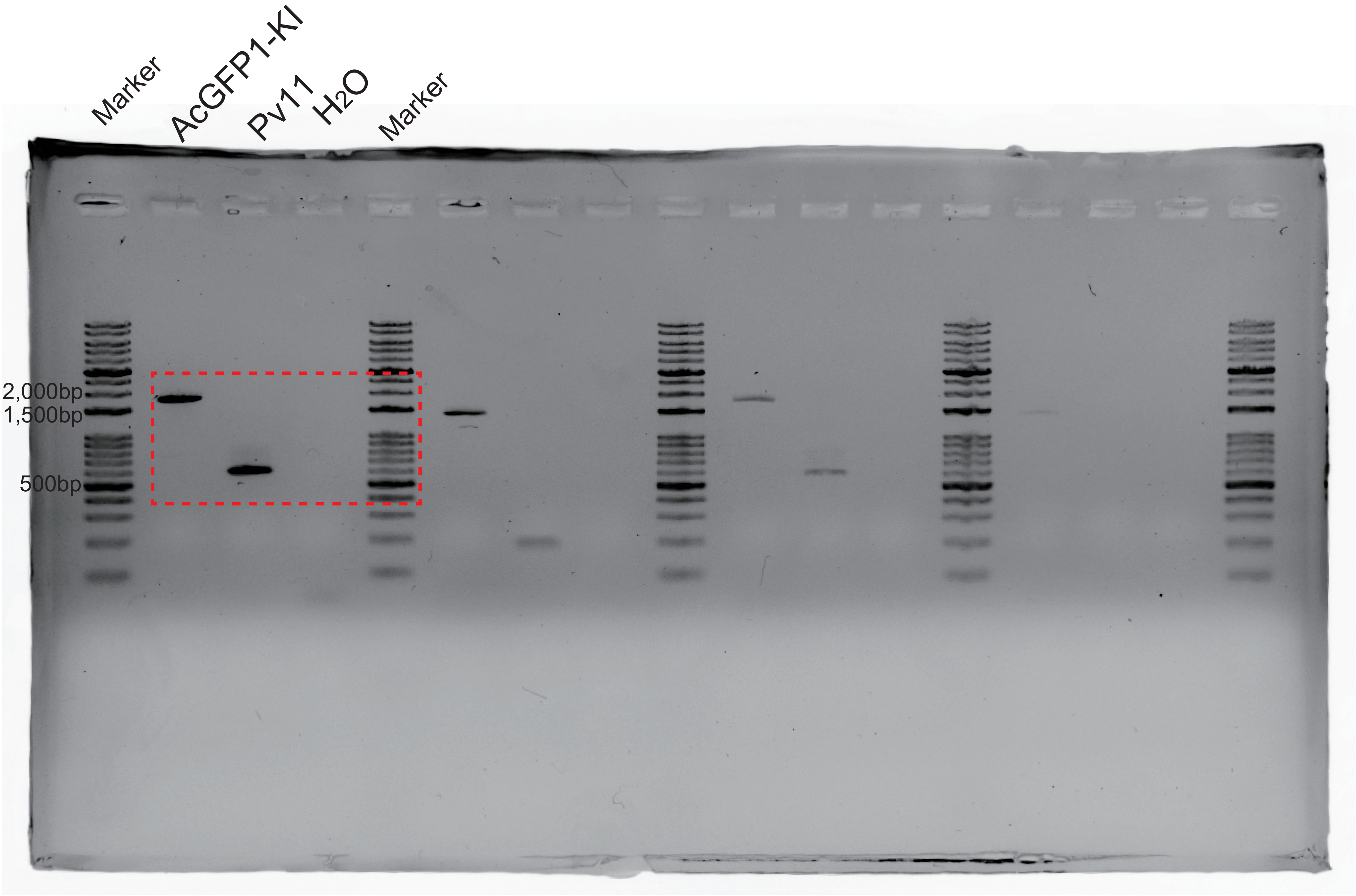

Supplementary Figure S3

Original PCR image in Figure 1c.

The cropped region in Figure 1c is enclosed in red dash line.

Fig. S4

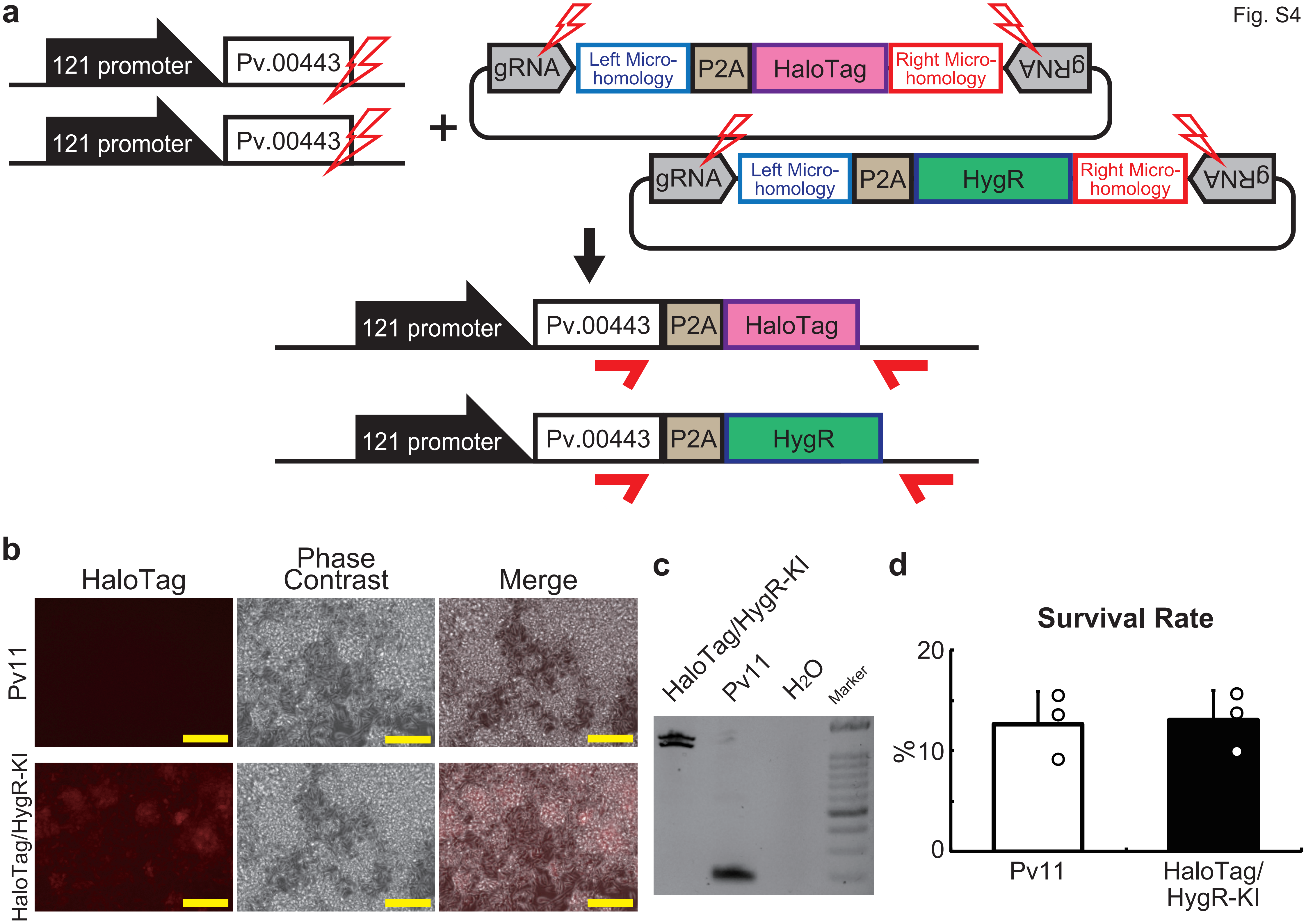
a
121 promoter
Pv.00443
121 promoter
Pv.00443
+
gRNA
Left Micro-homology
P2A
HaloTag
Right Micro-homology
gRNA
gRNA
Left Micro-homology
P2A
HygR
Right Micro-homology
gRNA
121 promoter
Pv.00443
P2A
HaloTag
121 promoter
Pv.00443
P2A
HygR
b
HaloTag
Phase Contrast
Merge
Pv11
HaloTag/HygR-KI
c
HaloTag/HygR-KI
Pv11
H2O
Marker
d
Survival Rate
20
%10
0
Pv11
HaloTag/HygR-KI

Supplementary Figure 4

The HaloTag or hygromycin resistance (HygR) gene was knocked in into 5' flanking site of the stop codon of *Pv.00443* gene.

**a)** The scheme of PITCh for HaloTag- or HygR-knock-in is shown; the Cas9- and gRNA-expression vectors and the donor vector harboring HaloTag or HygR gene was transfected into Pv11 cells. Red arrows indicate the primers used in genomic PCR for either intact Pv11 cells or the HaloTag/HygR- KI cells after selection by hygromycin treatment and sorting with fluorescence of HaloTag Ligand. **b)** The images of the cells were acquired by a microscope. **c)** Genomic PCR in the cell line was carried out. **d)** The survival rate after desiccation-rehydration treatment was compared between Pv11 and HaloTag-/HygR-KI cells. Scale bars, 100 μm. The values are expressed as mean ± SD; n = 3 in each group.

Fig. S5

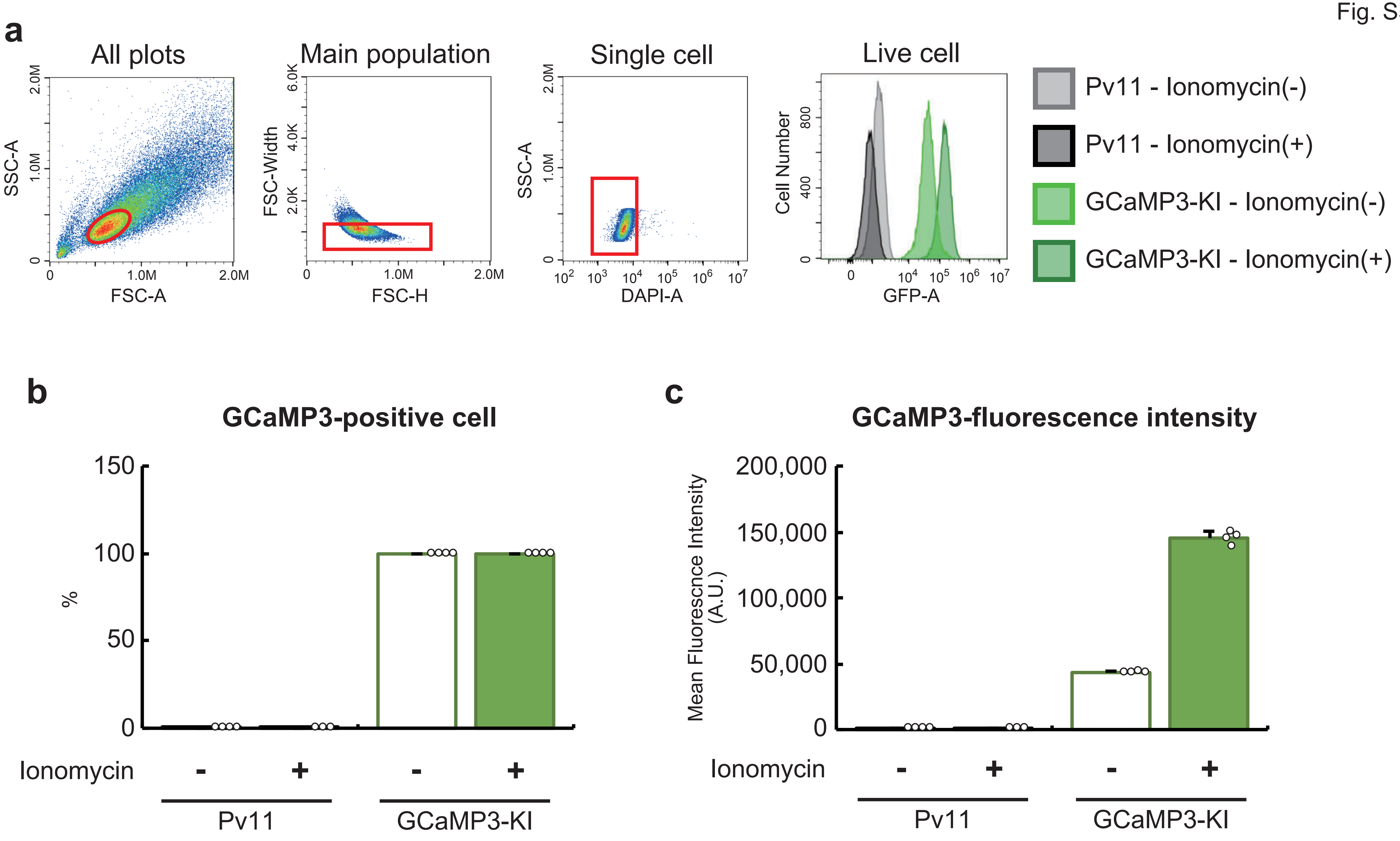
a
All plots
SSC-A
FSC-A
0
1.0M
2.0M
Main population
FSC-Width
FSC-H
2.0K
4.0K
6.0K
Single cell
SSC-A
DAPI-A
Live cell
Cell Number
GFP-A
400
800
Pv11 - Ionomycin(-)
Pv11 - Ionomycin(+)
GCaMP3-KI - Ionomycin(-)
GCaMP3-KI - Ionomycin(+)
b
GCaMP3-positive cell
%
0
50
100
150
Ionomycin
-
+
Pv11
GCaMP3-KI
c
GCaMP3-fluorescence intensity
Mean Fluorescnce Intensity (A.U.)
0
50,000
100,000
150,000
200,000
Ionomycin
-
+
Pv11
GCaMP3-KI

Supplementary Figure 5

Gating hierarchy and dot-plot images of GCaMP3-KI cells in flow cytometry experiments.

**a)** Representative dot-plot images are shown, and all gates are colored red. The gating hierarchy is main population – single cell – live cell – GCaMP3-positive cell. **b)** The proportions of GCaMP3-positive cells in the live cell population were analyzed. **c)** The mean fluorescent intensities in the live cell population were quantified. The values are expressed as mean ± SD; n = 3-4 in each group.

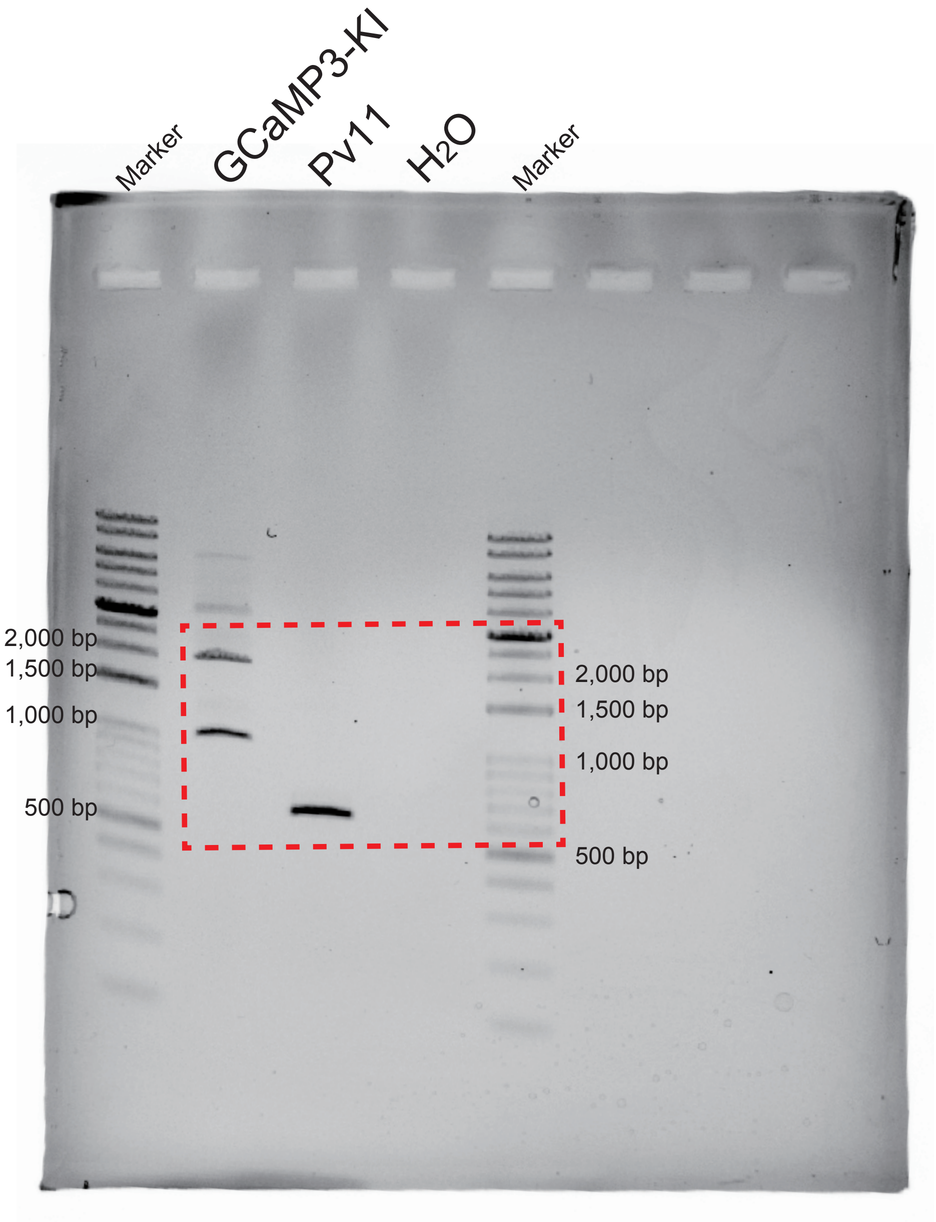
Marker
GCaMP3-KI
Pv11
H2O
Marker
2,000 bp
1,500 bp
1,000 bp
500 bp
2,000 bp
1,500 bp
1,000 bp
500 bp

Supplementary Figure 6

Original PCR image in Figure 2c.

The cropped region in Figure 2c is enclosed in red dash line.

Fig. S7

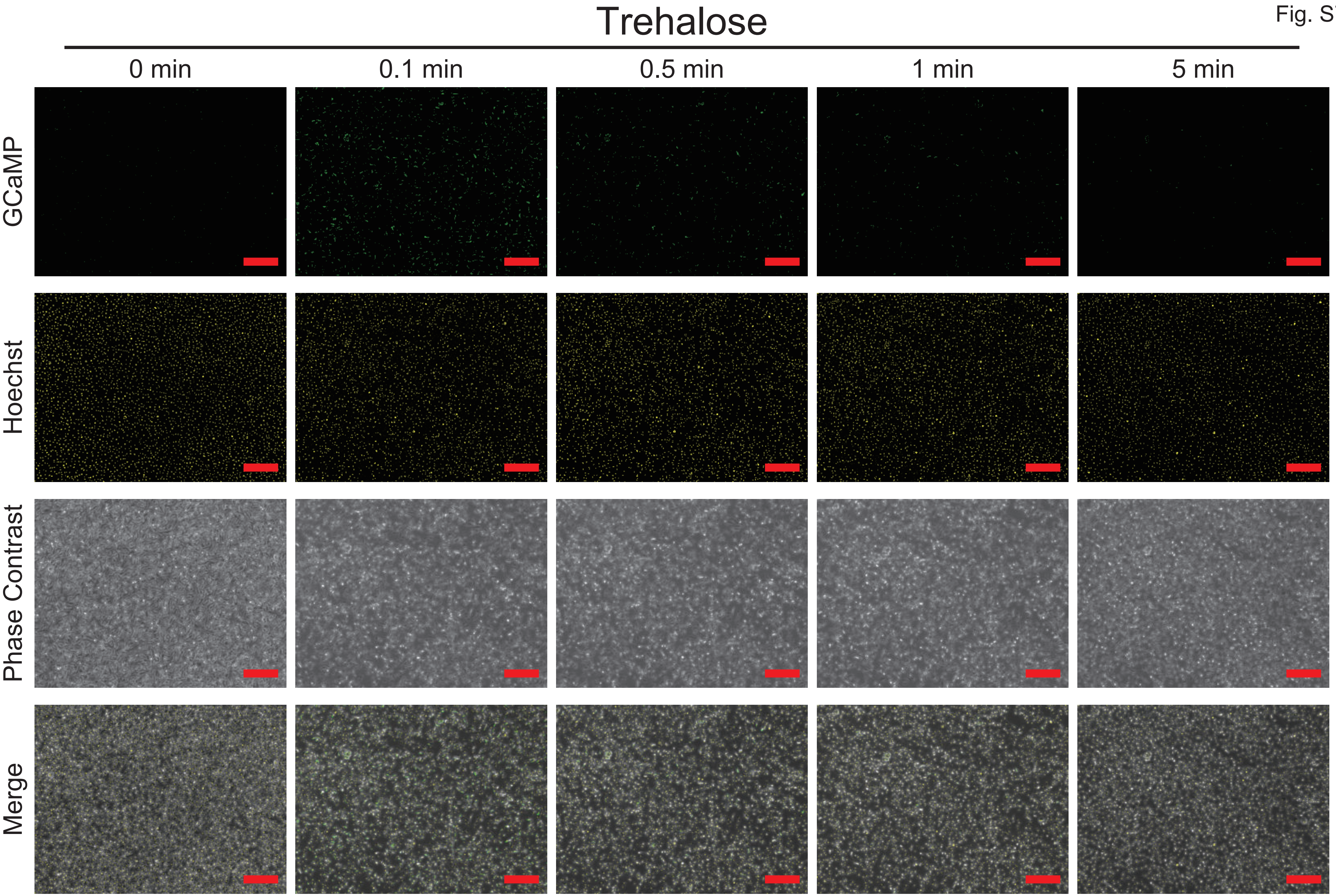

Trehalose
0 min
0.1 min
0.5 min
1 min
5 min
GCaMP
Hoechst
Phase Contrast
Merge

Supplementary Figure 7

Original images of “Trehalose” samples in Figure 3a. Scale bars, 100 μm.

Fig. S8

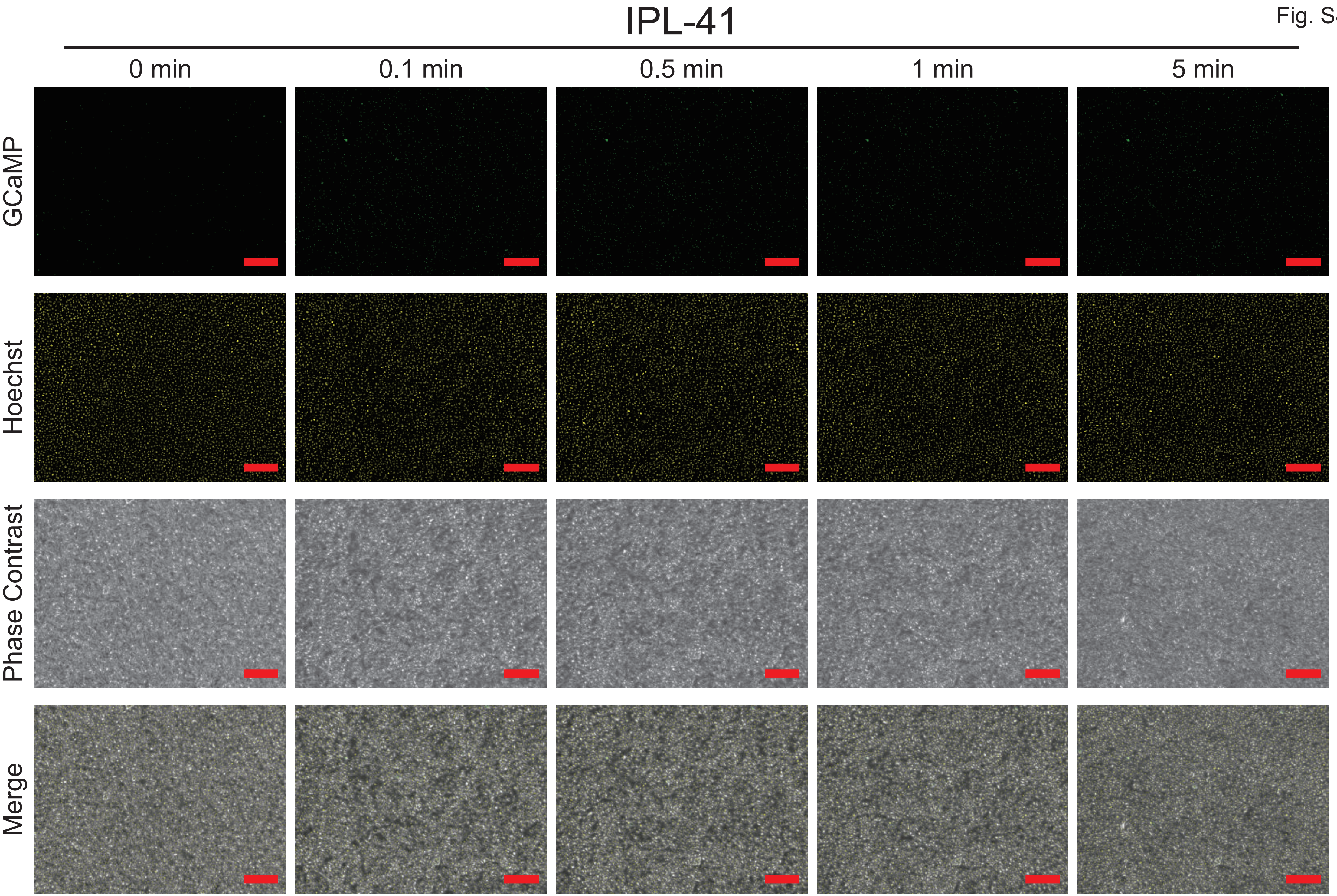
IPL-41
0 min
0.1 min
0.5 min
1 min
5 min
GCaMP
Hoechst
Phase Contrast
Merge

Supplementary Figure 8

Original images of “IPL-41” samples in Figure 3a. Scale bars, 100 μm.

Fig. S9

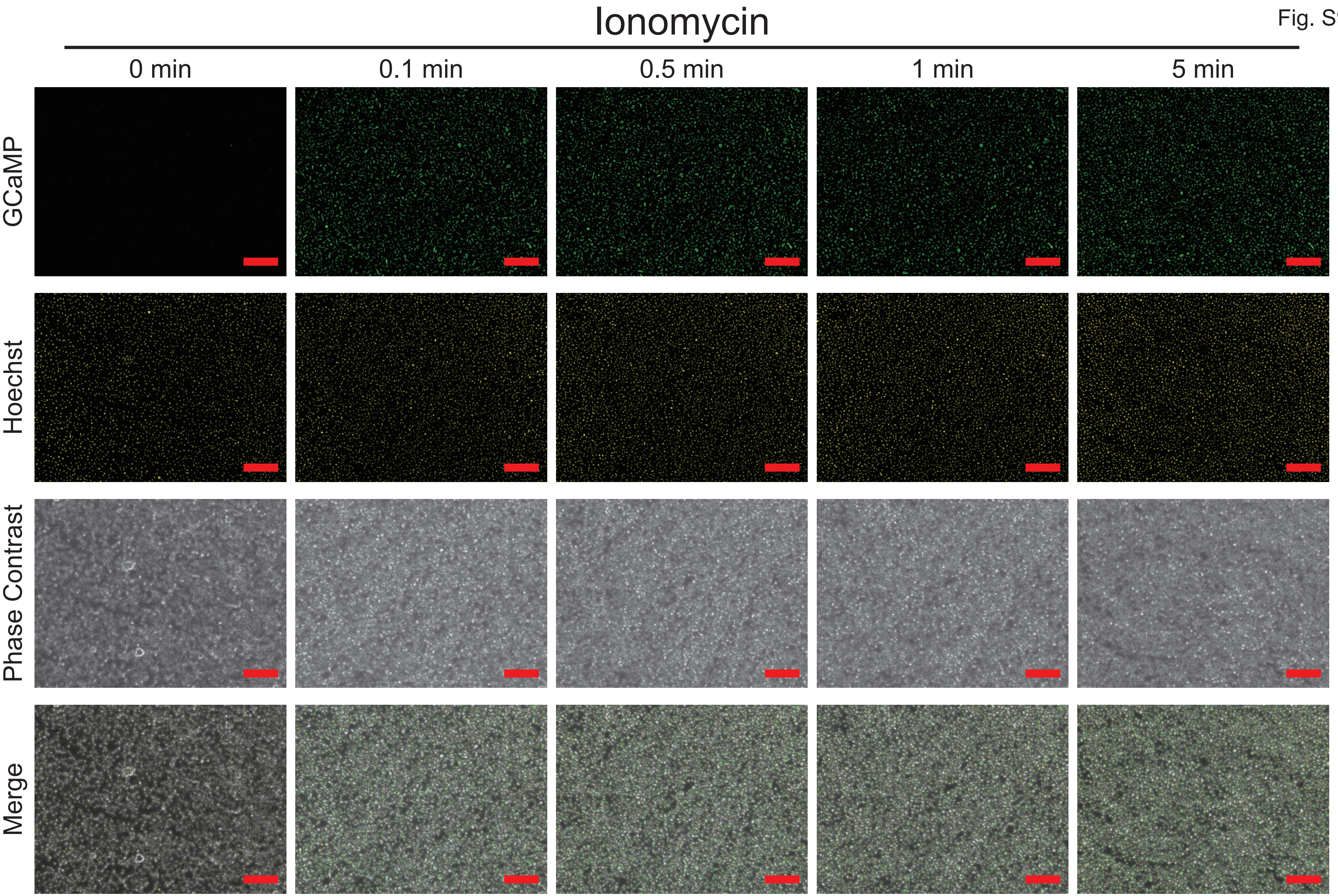
Ionomycin
0 min
0.1 min
0.5 min
1 min
5 min
GCaMP
Hoechst
Phase Contrast
Merge

Supplementary Figure 9
Ionomycin treatment in GCaMP3-KI cells.
GCaMP3-KI cells were treated with ionomycin, and the time-course images were acquired.
Scale bars, 100 μm.

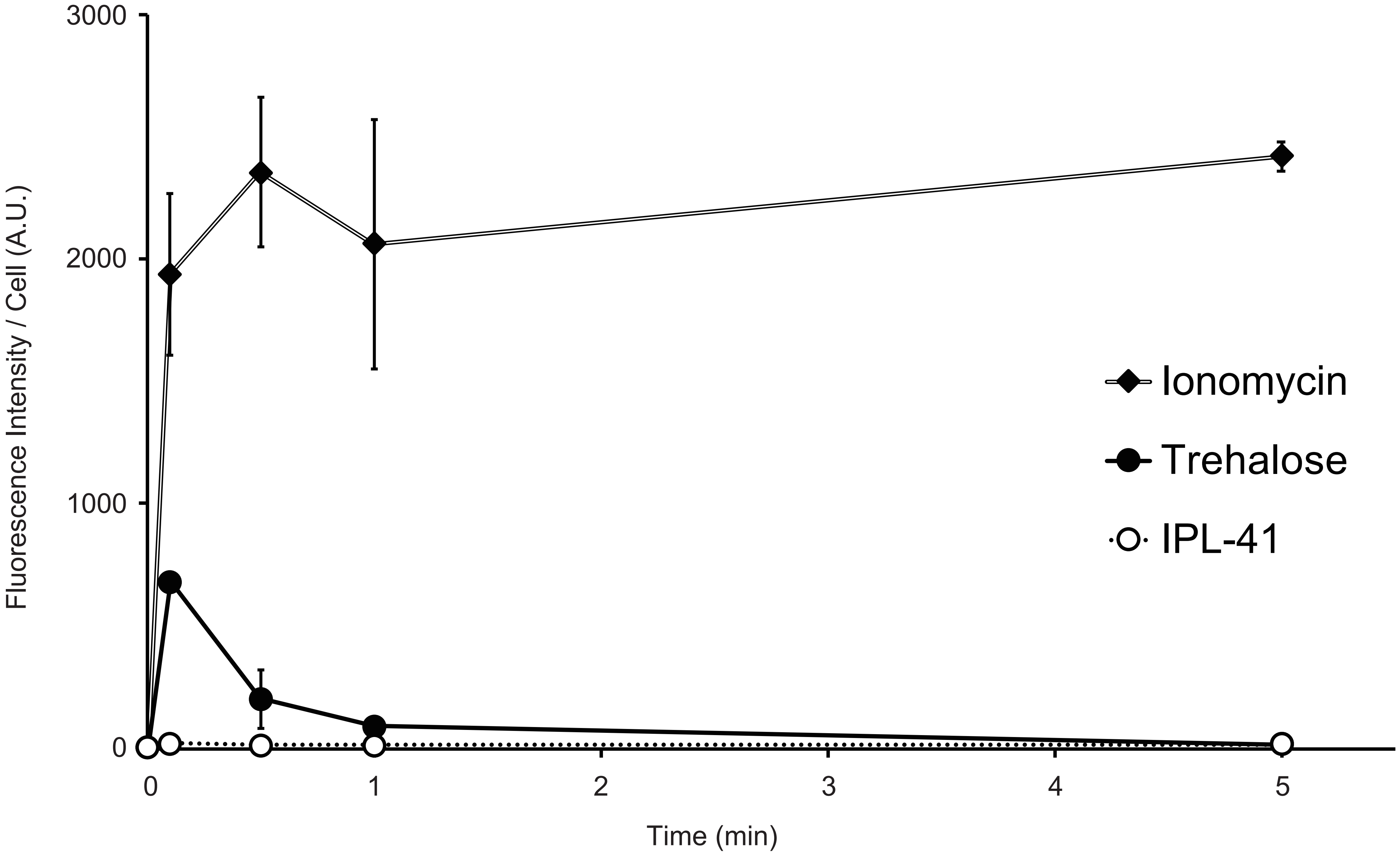
3000
2000
1000
0
Fluorescence Intensity / Cell (A.U.)
0
1
2
3
4
5
Time (min)
Ionomycin
Trehalose
IPL-41

Supplementary Figure 10

Quantitative data of GCaMP3-fluorescence with ionomycin-treated GCaMP3-KI cells. The values are expressed as mean ± SD; n = 3 in each group.

Fig. S11

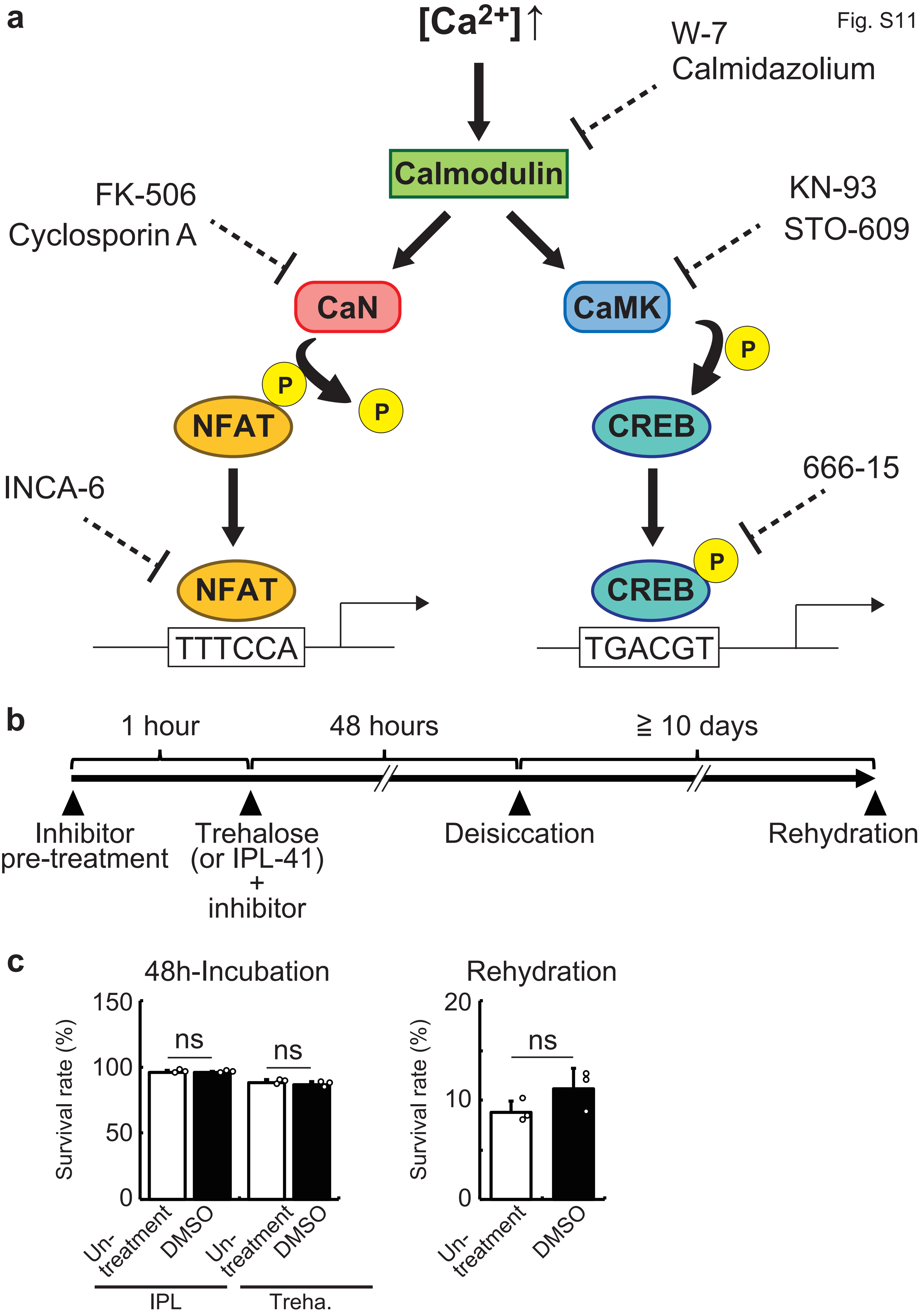
a
[Ca2+]↑
W-7
Calmidazolium
Calmodulin
FK-506
Cyclosporin A
KN-93
STO-609
CaN
CaMK
P
NFAT
CREB
INCA-6
666-15
TTTCCA
TGACGT
b
1 hour
48 hours
≧ 10 days
Inhibitor pre-treatment
Trehalose (or IPL-41) + inhibitor
Deisiccation
Rehydration
c
48h-Incubation
Rehydration
Survival rate (%)
150
100
50
0
20
10
ns
Un-treatment
DMSO
IPL
Treha.

Supplementary Figure 11

Contribution of the calcium signaling pathways to the desiccation tolerance

**a)** The representative calcium signaling pathways and the inhibitors used in this study were shown. **b)** The experimental scheme of the inhibitor experiments is shown. **c)** The effect of DMSO on the survival rates during IPL-41/trehalose incubation or after desiccation-rehydration treatment were analyzed. The values are expressed as mean ± SD; n = 3 in each group.

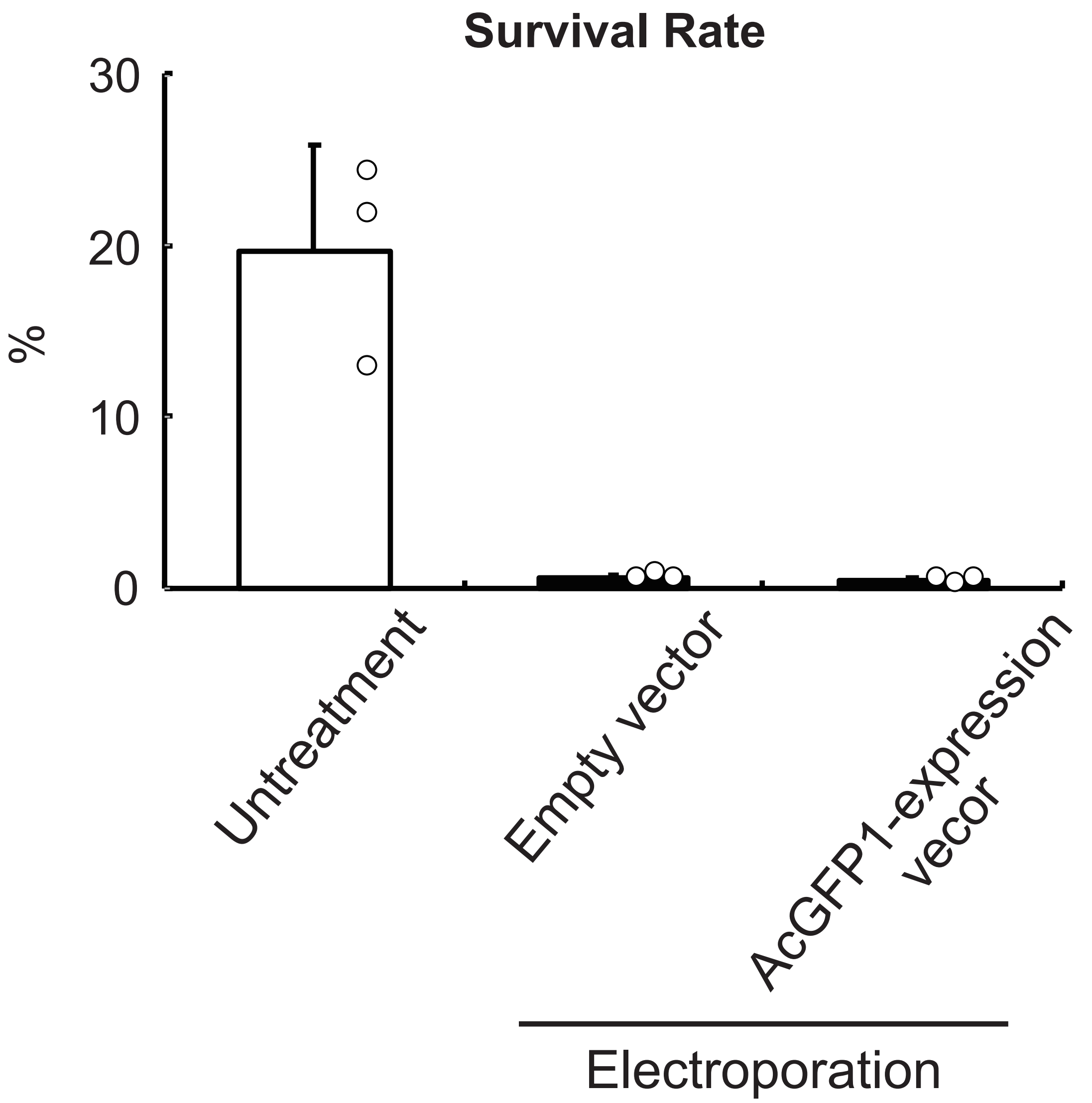
Survival Rate
30
20
10
0
%
Untreatment
Empty vector
AcGFP1-expression vecor
Electroporation

Supplementary Figure 12

Deleterious effect of electroporation on desiccation tolerance in Pv11 cells.

Electroporation was performed two days prior to trehalose treatment and the survival rate after desiccation-rehydration treatment were analyzed.
