## Supplementary material for "Cas9-mediated Genome Editing Reveals a Significant Contribution of Calcium Signaling Pathways to Anhydrobiosis in Pv11": Description of Supplementary Data

Description of Additional Supplementary Files

Supplementary Data 1

The full-length sequence of pPvU6b-DmtRNA-BbsI plasmid

Supplementary Data 2

The full-length sequence of the donor vector, pCRII-Pv.00443#1µH-P2A-AcGFP1-P2A-ZeoR

Supplementary Data 3

The genome sequence of AcGFP1-KI cell.

Supplementary Data 4

The genome sequence of HaloTag-knock-in allele.

Supplementary Data 5

The genome sequence of BlaR-knock-in allele.

Supplementary Data 6

The genome sequence of GCaMP3-knock-in allele.

Supplementary Data 7

The genome sequence of ZeoR-knock-in allele.

Supplementary Data 8

The full-length sequence of pPvU6b-DmtRNA-AcGFP1#1 plasmid.

Supplementary Data 9

The full-length sequence of pPvU6b-DmtRNA-AcGFP1#3 plasmid.

Supplementary Data 10

The full-length sequence of pPvU6b-DmtRNA-Pv.00443#1 plasmid.

Supplementary Data 11

The full-length sequence of the donor vector, pCR4-Pv.00443#1µH-P2A-BbsI.

Supplementary Data 12

The full-length sequence of the donor vector, pCR4-Pv.00443#1µH-P2A-BlaR.

Supplementary Data 13

The full-length sequence of the donor vector, pCR4-Pv.00443#1µH-P2A-BlaR.

Supplementary Data 14

The full-length sequence of the donor vector, pCR4-Pv.00443#1µH-P2A-GCaMP3.

Supplementary Data 15

The full-length sequence of the donor vector, pCR4-Pv.00443#1µH-P2A-ZeoR.

Supplementary Data 16

Source data underlying plots shown in the figures.
